# The thresholding problem and variability in the EEG graph network parameters

**DOI:** 10.1101/2022.01.26.477863

**Authors:** Timofey Adamovich, Ilya Zakharov, Anna Tabueva, Sergey Malykh

## Abstract

Graph thresholding is a frequently used practice of eliminating the weak connections in brain functional connectivity graphs. The main aim of the procedure is to delete the spurious connections in the data. However, the choice of the threshold is arbitrary, and the effect of the threshold choice is not fully understood. Here we present the description of the changes in the global measures of a functional connectivity graph depending on the different proportional thresholds based on the 146 resting-state EEG recordings. The dynamics is presented in five different synchronization measures (wPLI, ImCoh, Coherence, ciPLV, PPC) in sensors and source spaces. The analysis shows significant changes in the graph’s global connectivity measures as a function of the chosen threshold which may influence the outcome of the study. The choice of the threshold could lead to different study conclusions; thus it is necessary to improve the reasoning behind the choice of the different analytic options and consider the adoption of different analytic approaches. We also proposed some ways of improving the procedure of thresholding in functional connectivity research.

## Introduction

Brain connectomics is a rapidly developing field of research that investigates the interactions between relatively homogeneous brain cell circuits (or brain areas). In response to the external environment, the various areas across the cortex begin to interact, forming a macroscale whole-brain network [1]. One way to understand this network comes from mathematical graph theory which can describe complex structures and dynamics within networks [2]. The network approach is based on a pairwise matrix of connections between nodes and was used in various studies, to examine the relations between brain characteristics of brain networks with different psychological phenomena, such as cognitive processes, personality, or psychopathology [3–8] based on functional magnetic resonance imaging (fMRI) or diffusion tensor imaging (DTI), electroencephalography (EEG), magnetoencephalography (MEG) and functional near-infrared spectroscopy (fNIRS) [9, 10] data.

Network neuroscience is a new and developing area, and one of its challenges is considerable variability in the connectivity calculation routines, which is not always detailed and explicitly motivated in studies. Estimation of the functional connections graph via various indexes (see [11] for a review) creates a full adjacency matrix that makes no sense anatomically or mathematically. The next step is usually the creation of the sparse matrix by choosing some threshold value as a cut-off point below which the values are removed [12]. However, while thresholding is a good way to reduce the possibility of too much influence of spurious or weak connections, it leads to confounds related to the arbitrary choice of the calculation routines.

There is no consensus in the scientific society as to how to choose the specified thresholds. There are two main ways to estimate the threshold value - an absolute and proportional thresholding. An absolute threshold is a specific value of a connection strength, and a proportional threshold is a percentage, below which the connection is considered spurious and deleted from the graph. An important issue is the selection of the specific value (both absolute and proportional) and, accordingly, the number of edges in the resulting network. The thresholding procedure has been criticized in numerous studies, both from theoretical and practical positions [2]. The choice of the threshold could lead to incomparable results [13, 14]. In a recent fMRI study, the range of absolute thresholds varied from 0.1 to 0.8 correlation coefficient and proportional thresholds from 2 to 40% leading the graphs to be incomparable [15, 16]. The fMRI graph-based connectivity metrics are unstable across various threshold values (and overall, more stable across proportional thresholds compared to absolute [13]). In a recent DTI study (Diffusion Tensor Tomography, [16]) the thresholding procedure has been shown to lead to differences in connectivity metrics if more than 70% of the connections between the nodes were removed. Also, it has been shown that calculating the connectivity metrics without thresholding is feasible and, thus, there is now no practical utility in this step of the analysis for DTI data.

While the thresholding procedure is widely used, there is also another, data-driven way to prepare brain data for network analysis, which is based on an algorithmic reduction of the number of nodes. The most popular algorithms are based on the spanning tree procedure (e.g. minimum or condensed spanning tree MST or CST, respectively), which constructs a unique and acyclic sub-network with a fixed number of connections (see, for example, [17, 18]). In the brain imaging data, the spanning-tree approach could be very useful, but a direct comparison of the results obtained with thresholding approaches and various data-driven approaches is needed.

Initially, connectivity studies were mostly based on various types of MRI (both structural and functional) data. However, a considerable part of brain activity cannot be captured by MRI due to its poor temporal resolution. The M/EEG allows the analysis of quick or high-frequency oscillations [11, 19] and reflects the efficiency of communication between brain areas [19, 20]. But, according to the recent International Federation of Clinical Neurophysiology (IFCN) report, for M/EEG studies “there is no clear consensus on the statistical thresholds for the confirmation of a significant effect at the scalp or source-level” [21]. As far as we know the effect of the thresholding for the calculation of the EEG-derived connectivity graph metrics has never been directly assessed.

The article is based on a study first reported in the Journal of Physics: Conference Series [22] and presents an extended analysis.

## Results

### The distribution of the connectivity strength

The connectivity graphs could be influenced by the individual differences in connectivity strength. Figure 1 illustrates the descriptive statistics of individual connectivity strength distributions. The plots are presented for each connectivity measure in the sensor (left column) and source (right column) space. From top to bottom: Median, Mean, Median Absolute Deviation, SD.

**Figure 1.**
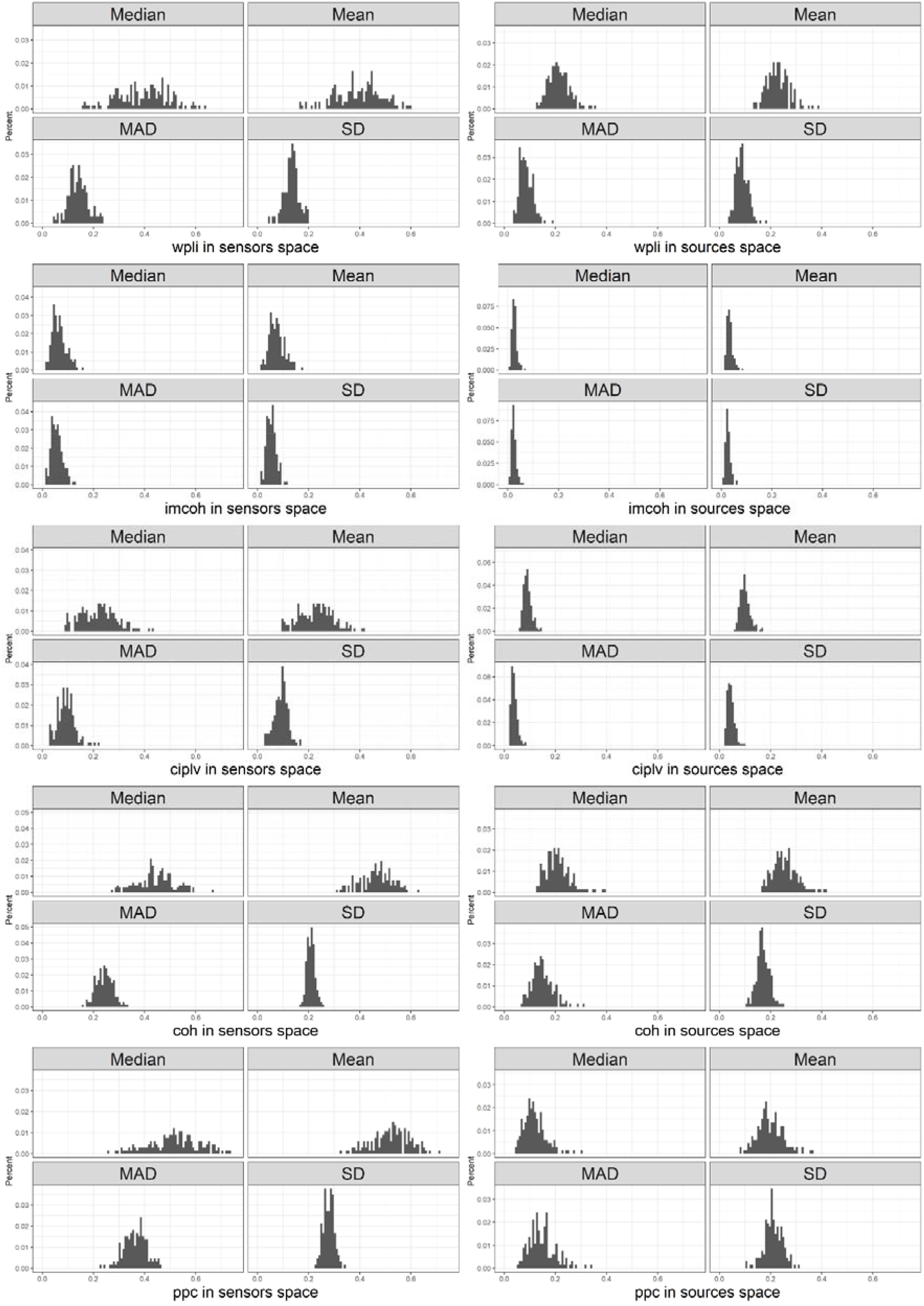
Distribution of descriptives for connectivity measures in sensor and source space.

The shape of the distribution of the mean and standard deviation in the sample point out the variability of absolute values and fluctuations in the shape of the individual connectivity strength distributions. The differences in the absolute connectivity strength could impact the graph measures (mostly connection-weight based, e.g. path length, [14], however, structural properties of the graph should remain the same. For the thresholds that lead to low graph densities the problem of a disconnected graph might appear which will lead to the inability to estimate the path length and clustering coefficient of the overall network.

To overcome the disconnected graphs problem the procedure of eliminating the isolated nodes from the graph is often used. Isolated nodes almost inevitably appear with high thresholds, thus leading to the differences in the number of nodes in the graphs which could lead to biased results. According to our data (see Appendix) the number of isolated nodes increases with a higher threshold level to the point where there is no matrix without a disconnected node. It is possible that on the higher thresholds graph measures are derived from the graph with different sets of nodes. A thorough discussion of the missing nodes is beyond the scope of the paper.

### Threshold-level induced changes in the graph measures

In the next step of our analysis to investigate the effect of the thresholding values on the chosen graph metrics we trace the changes in the graph measures retrieved from the different chosen thresholds. The results are presented in figures 2-3.

**Figure 2.**
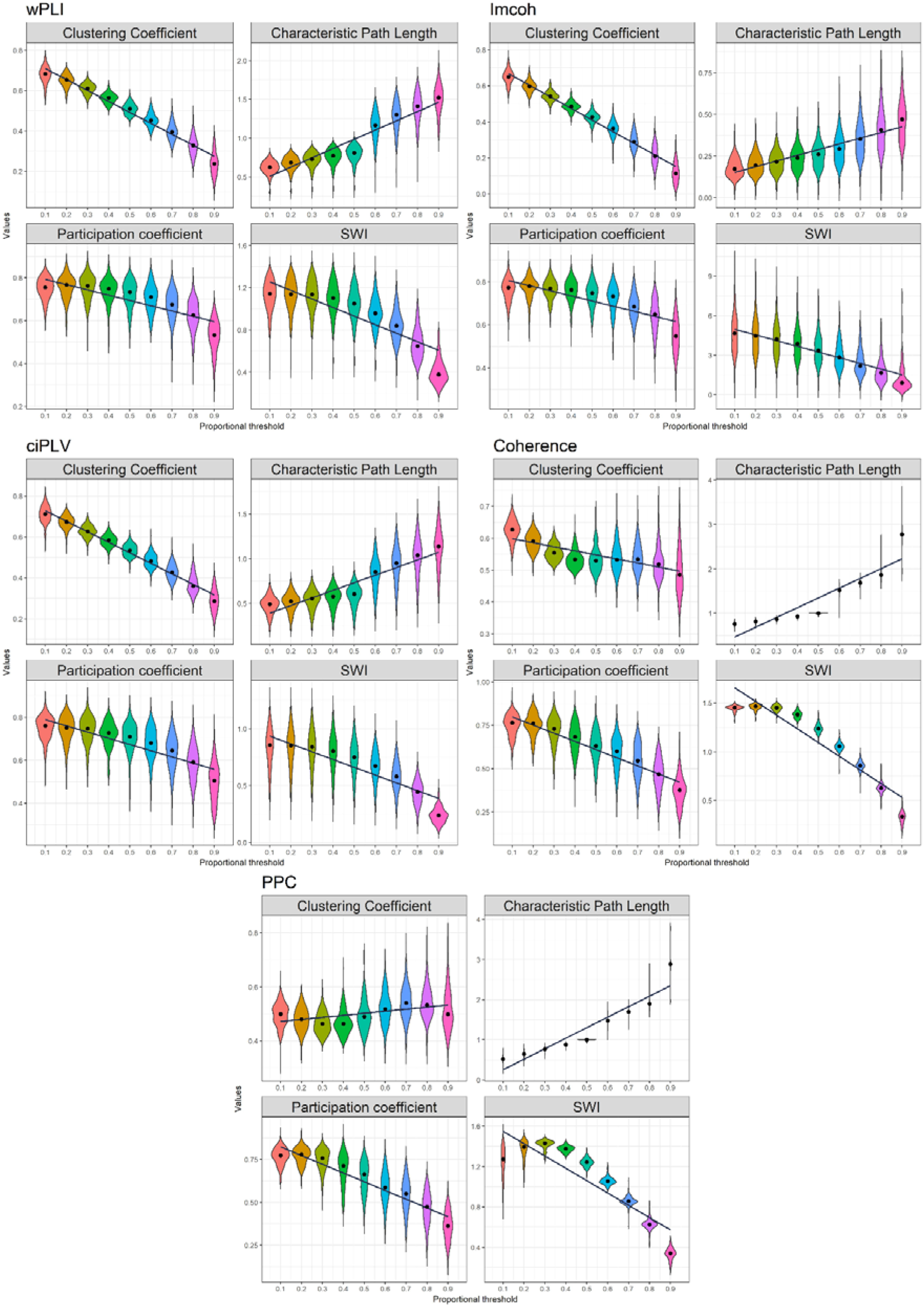
Effect of density thresholding on graph measures in sensor space. Five measures are presented in the following order (top-to-bottom): wPLI, imcoh, ciPLV, coherence, PPC On each plot four measures are presented (left-to-right, top-to-bottom): Clustering Coefficient, Characteristic PL, Participation Coefficient, SWI. The black line represents linear regression over the data.

**Figure 3.**
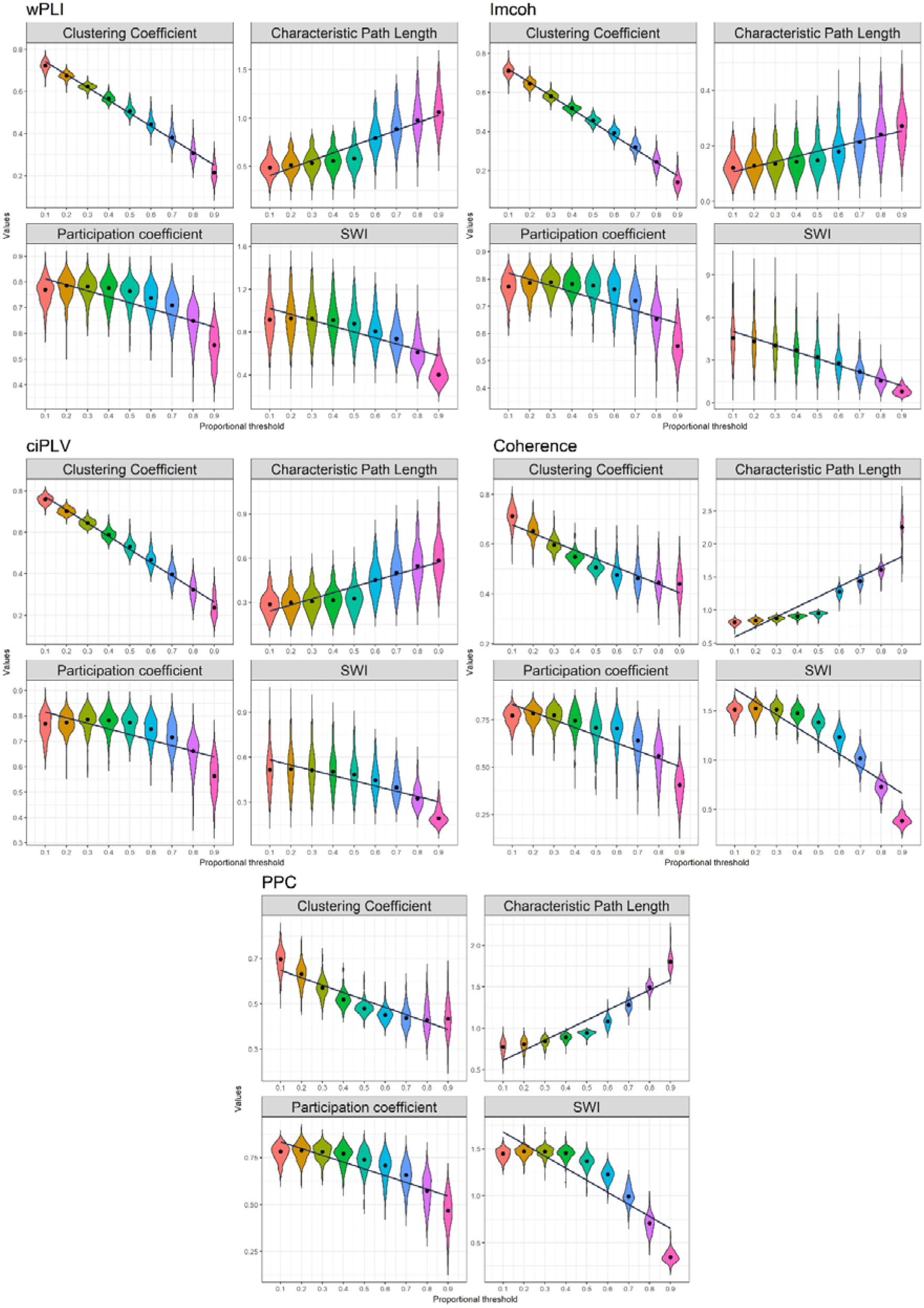
Effect of density thresholding on graph measures in source space. Five measures are presented in the following order (left-to-right, top-to-bottom): wPLI, imcoh, ciPLV, coherence, PPC. On each plot, four measures are presented (left-to-right, top-to-bottom): Clustering Coefficient, Characteristic PL, Participation Coefficient, SWI. The black line represents linear regression over the data.

Characteristic Path length increases with the decreasing density, Clustering coefficient, Participation coefficient, and SWI decrease with decreasing density. Also, there are differences in the variance on the different thresholds with higher variance on the higher thresholds. The dynamics of the differences are similar in all synchronization measures and both spaces.

### Linear regression analysis of the threshold-level induced dynamics

At the next step of our analysis, we wanted to estimate the proportion of the variance in the calculation of graph metrics as the result of the thresholding procedure. Linear regression analysis was performed to investigate the dependence of measures on the thresholding value. The results are presented in Table 1. All coefficients are significant (p<0.05)

**Table 1.**
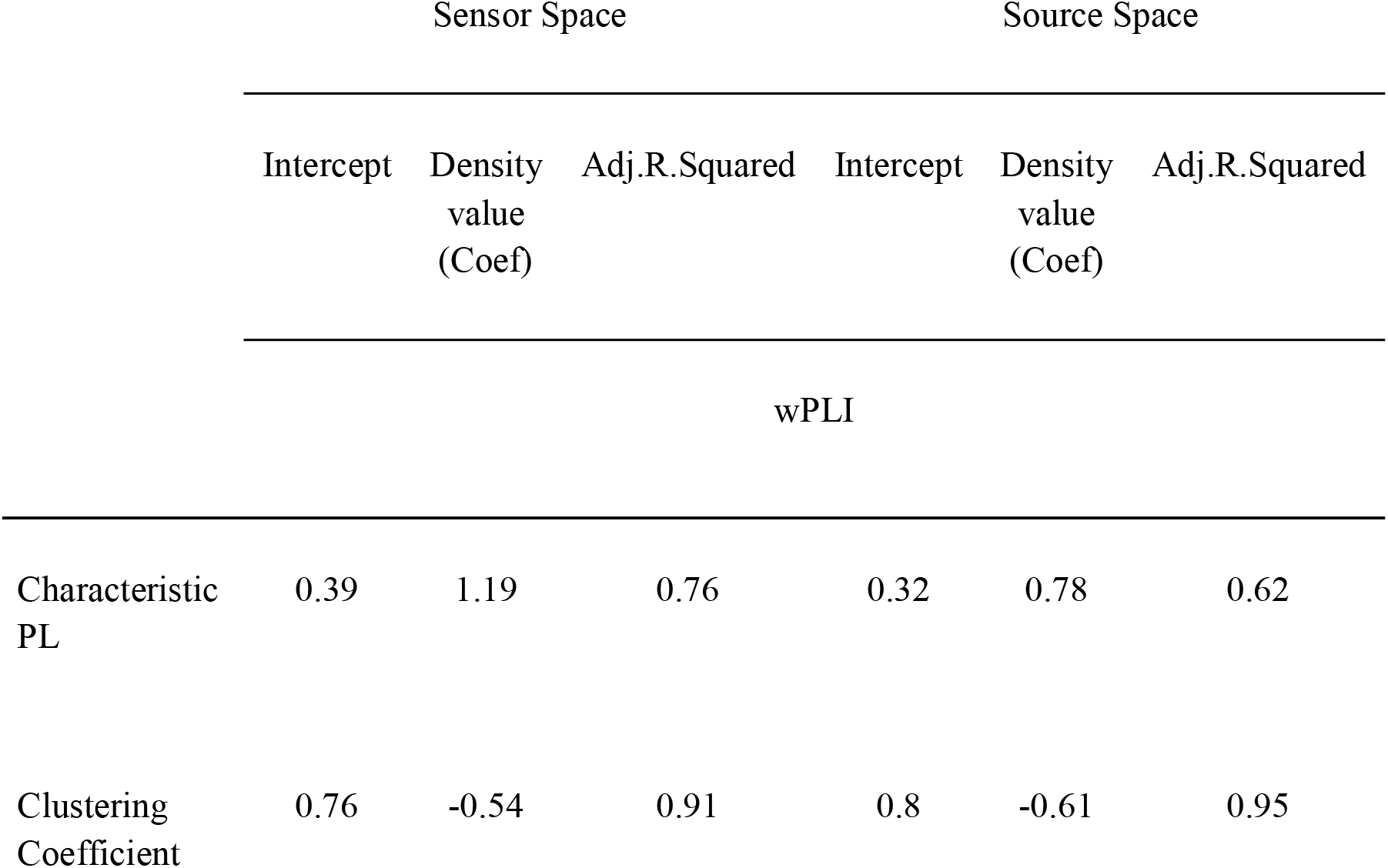
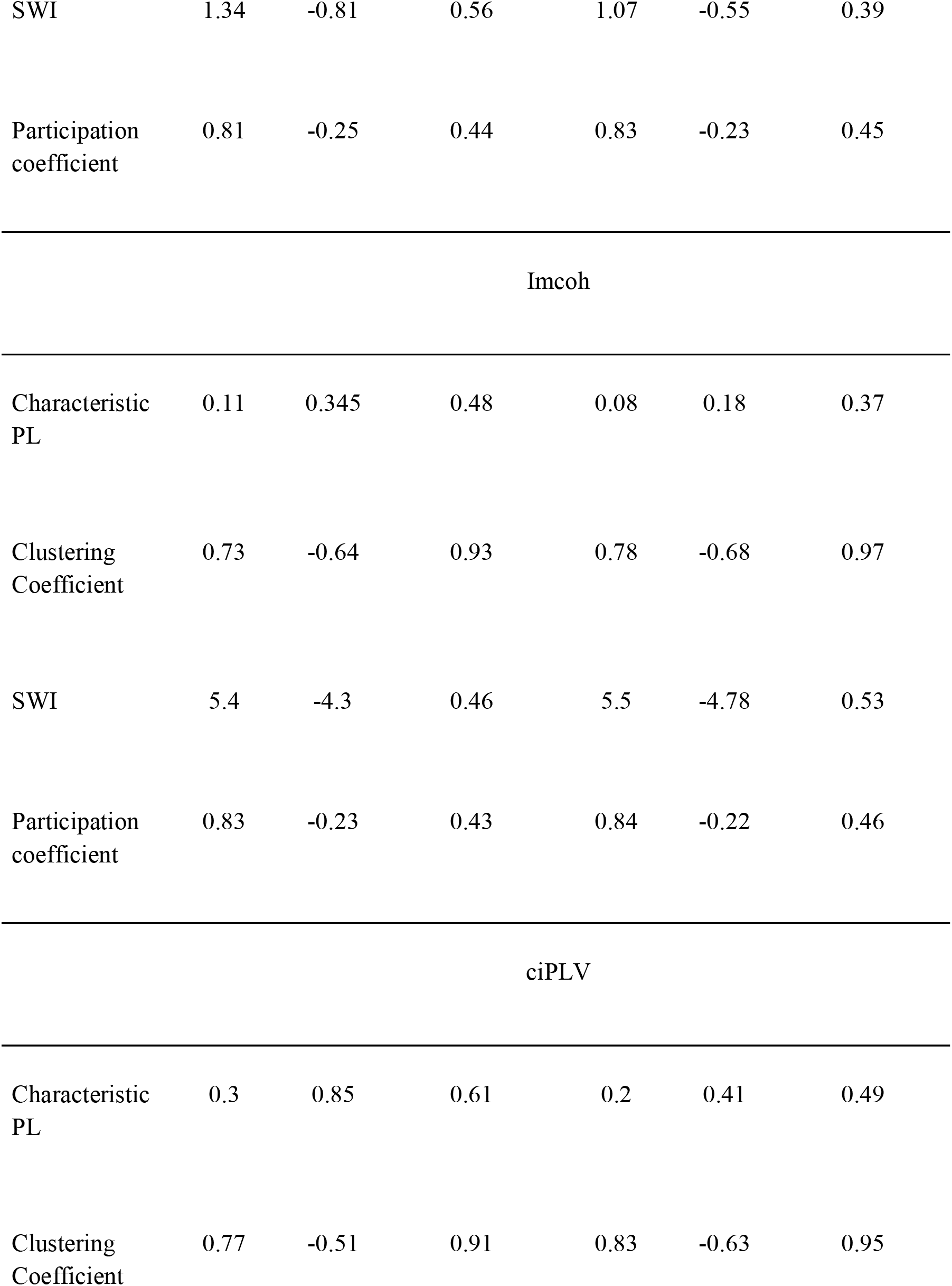
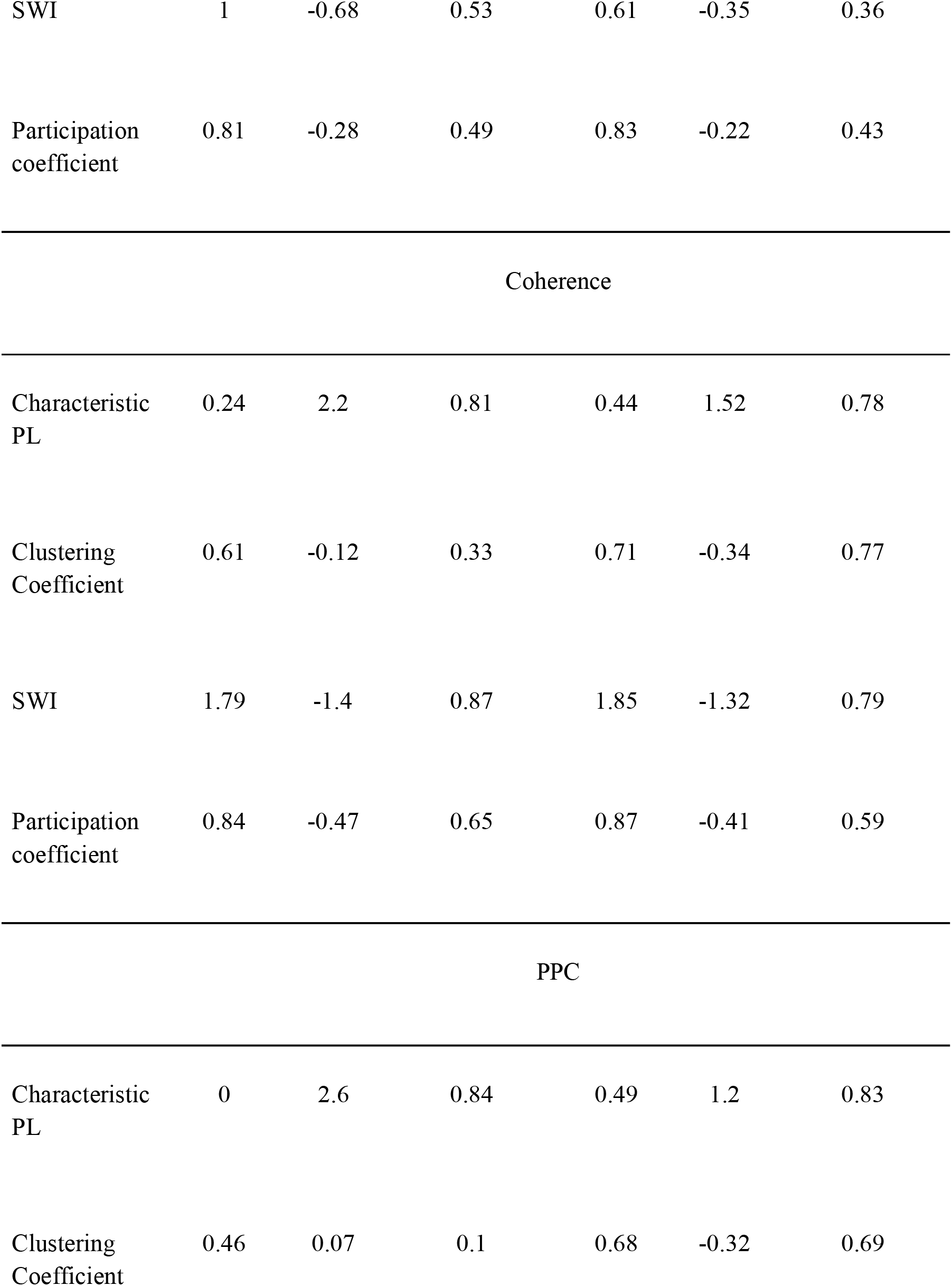
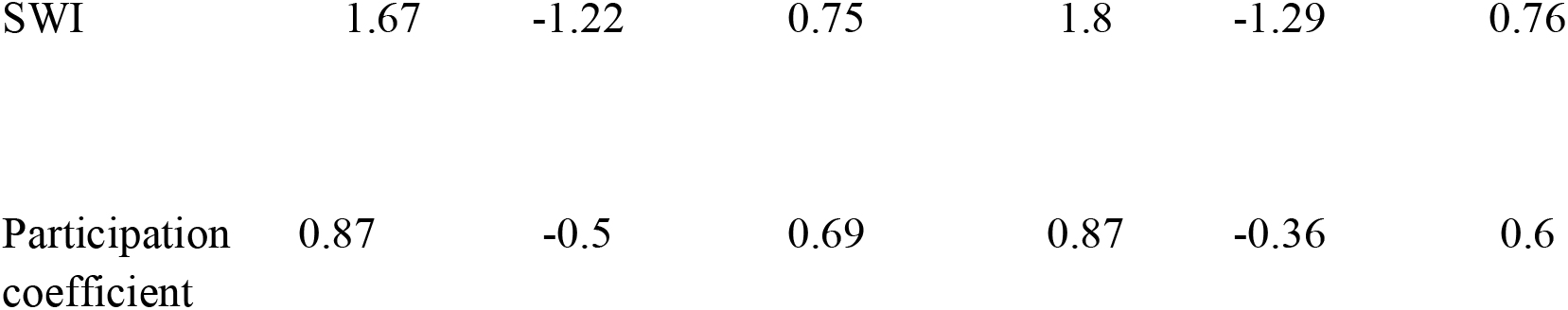
Linear regression of the brain connectivity measures.

|  | Sensor Space |  |  | Source Space |  |  |
| --- | --- | --- | --- | --- | --- | --- |
|  | Intercept | Density<br>value<br>(Coef) | Adj.R.Squared | Intercept | Density<br>value<br>(Coef) | Adj.R.Squared |
|  | wPLI |  |  |  |  |  |
| Characteristic<br>PL | 0.39 | 1.19 | 0.76 | 0.32 | 0.78 | 0.62 |
| Clustering<br>Coefficient | 0.76 | -0.54 | 0.91 | 0.8 | -0.61 | 0.95 |
| SWI | 1.34 | -0.81 | 0.56 | 1.07 | -0.55 | 0.39 |
| Participation coefficient | 0.81 | -0.25 | 0.44 | 0.83 | -0.23 | 0.45 |
| Characteristic PL | 0.11 | 0.345 | 0.48 | 0.08 | 0.18 | 0.37 |
| Clustering Coefficient | 0.73 | -0.64 | 0.93 | 0.78 | -0.68 | 0.97 |
| SWI | 5.4 | -4.3 | 0.46 | 5.5 | -4.78 | 0.53 |
| Participation coefficient | 0.83 | -0.23 | 0.43 | 0.84 | -0.22 | 0.46 |
| Characteristic PL | 0.3 | 0.85 | 0.61 | 0.2 | 0.41 | 0.49 |
| Clustering Coefficient | 0.77 | -0.51 | 0.91 | 0.83 | -0.63 | 0.95 |
| SWI | 1 | -0.68 | 0.53 | 0.61 | -0.35 | 0.36 |
| Participation coefficient | 0.81 | -0.28 | 0.49 | 0.83 | -0.22 | 0.43 |
| Characteristic PL | 0.24 | 2.2 | 0.81 | 0.44 | 1.52 | 0.78 |
| Clustering Coefficient | 0.61 | -0.12 | 0.33 | 0.71 | -0.34 | 0.77 |
| SWI | 1.79 | -1.4 | 0.87 | 1.85 | -1.32 | 0.79 |
| Participation coefficient | 0.84 | -0.47 | 0.65 | 0.87 | -0.41 | 0.59 |
| Characteristic PL | 0 | 2.6 | 0.84 | 0.49 | 1.2 | 0.83 |
| Clustering Coefficient | 0.46 | 0.07 | 0.1 | 0.68 | -0.32 | 0.69 |
| SWI | 1.67 | -1.22 | 0.75 | 1.8 | -1.29 | 0.76 |
| Participation<br>coefficient | 0.87 | -0.5 | 0.69 | 0.87 | -0.36 | 0.6 |

There is a substantial linear trend in the dynamic of the measure changes, as reflected in the R-squared values. The R-squared ranges from 0.1 to 0.97 with a median value of 0.62. R-squared also varies in the different measures and spaces. Although there is a strong linear component in the measure’s dynamics, most of them show a more complex pattern of changes. At low thresholds, the changes are smaller than at the high thresholds. The clustering coefficient demonstrates an even more peculiar pattern of changes in some cases, it has a plateau in the middle thresholds.

### Correlational analysis of measures on the different thresholds

Correlations between the different thresholds were used to identify the relations between the different thresholds. In figure 4, we present the correlations between measures on different thresholds. On the top half, sensor-space correlations are presented, on the lower - source-space. Significant correlations (p>0.05) are shown in color.

**Figure 4.**
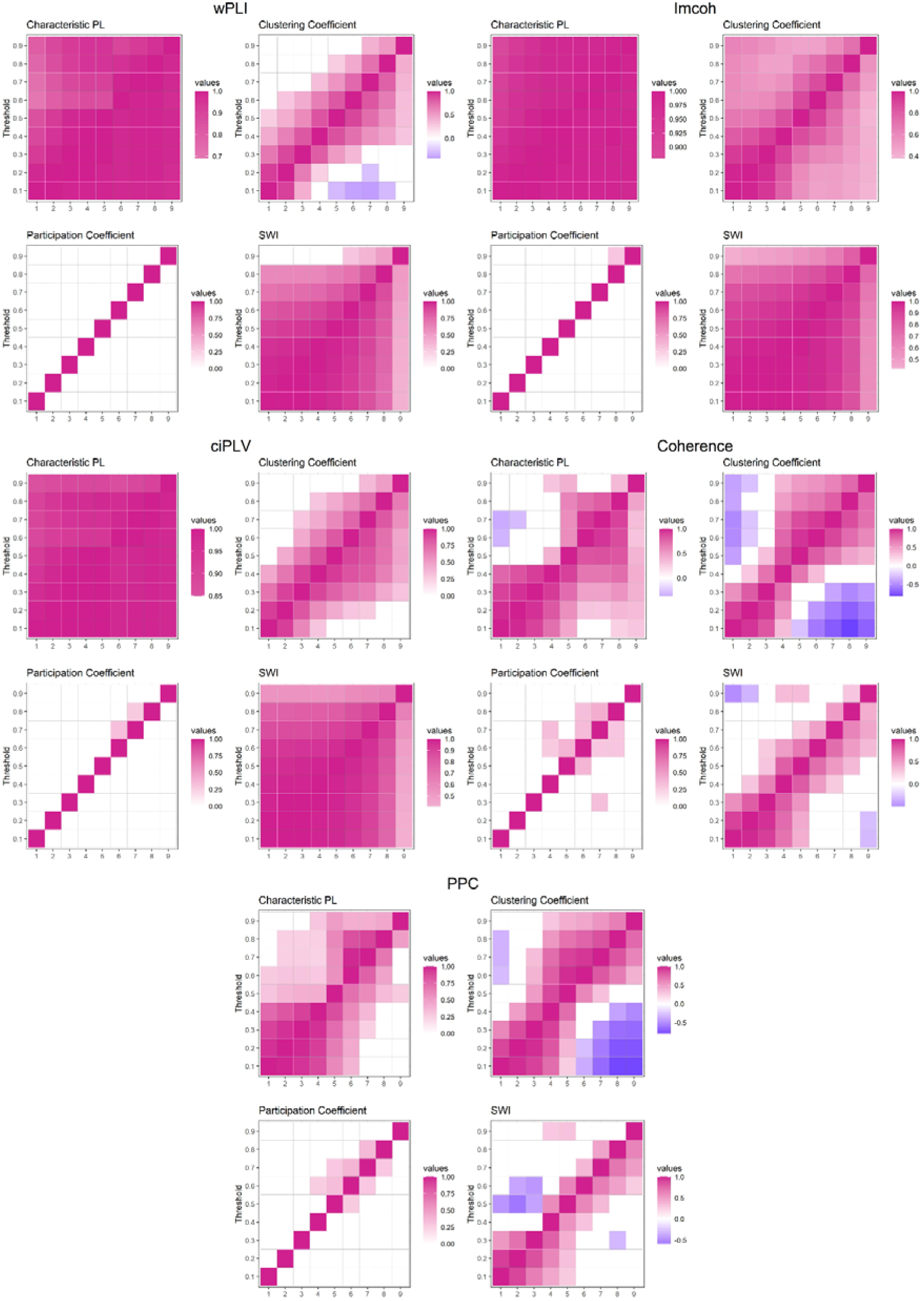
Correlation of graph measures on different thresholds. Five measures are presented in the following order (left-to-right, top-to-bottom): wPLI, imcoh, ciPLV, coherence, PPC. On each plot four measures are presented (left-to-right, top-to-bottom): Clustering Coefficient, Characteristic PL, Participation Coefficient, SWI. The utmost X-axis value on the right is the value obtained via OMST. Colored cells indicate the correlation between measures on two thresholds, purple color represents positive correlation, blue color represents negative correlation. White cells indicate absence of significant correlation. Transparency of color is related to the correlation strength, brighter color marks stronger correlations.

**Figure 5.**
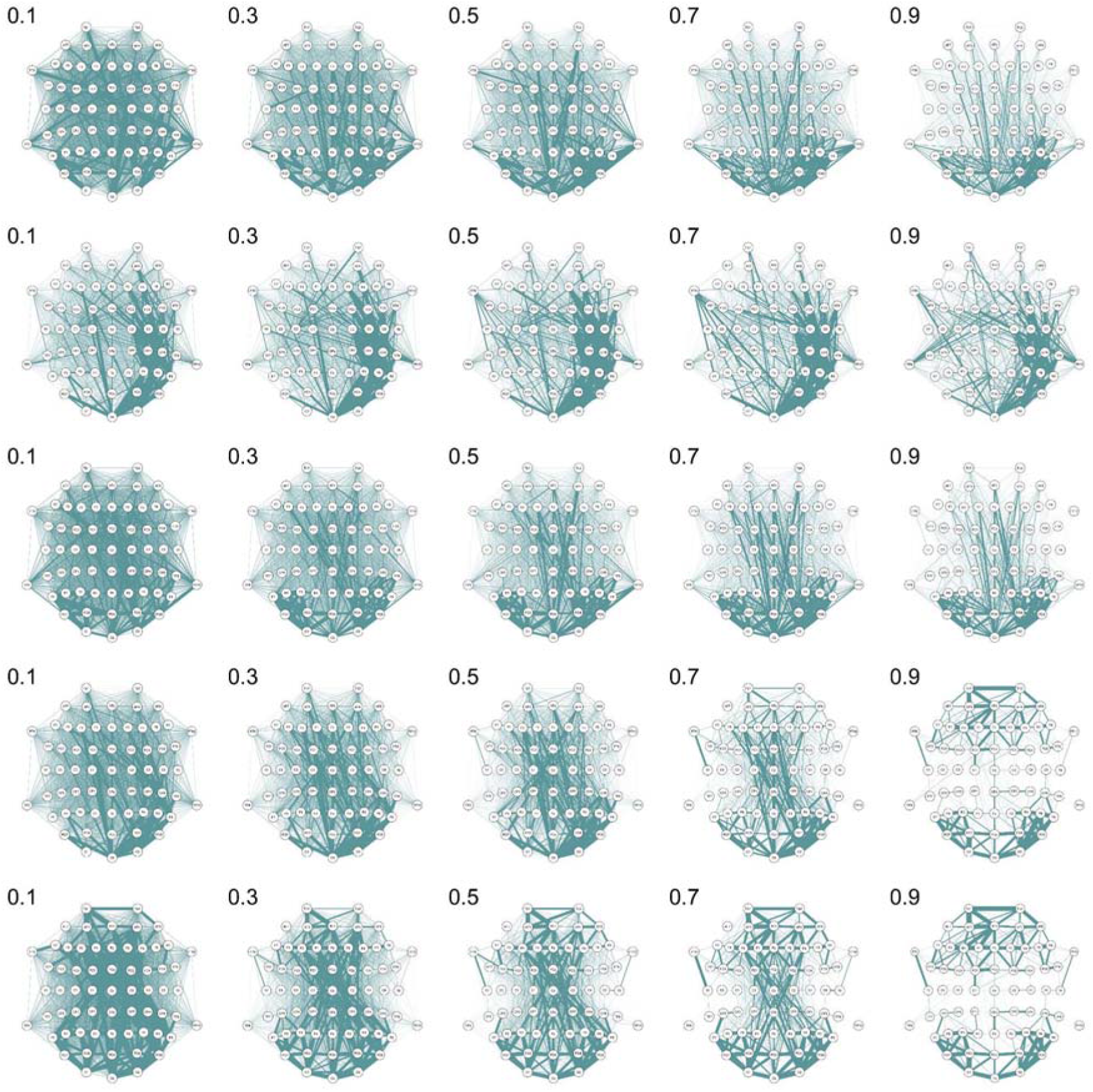
Edge probability graphs across different thresholds in sensor space. The thickness of edges reflects the relative probability of edge existence. Thicker edges are more frequent in the individual networks in the sample. The edge weights were scaled to the range [0, 1] across all thresholds. From top to bottom: wPLI, Imcoh, ciPLV, Coherence, PPC

**Figure 6.**
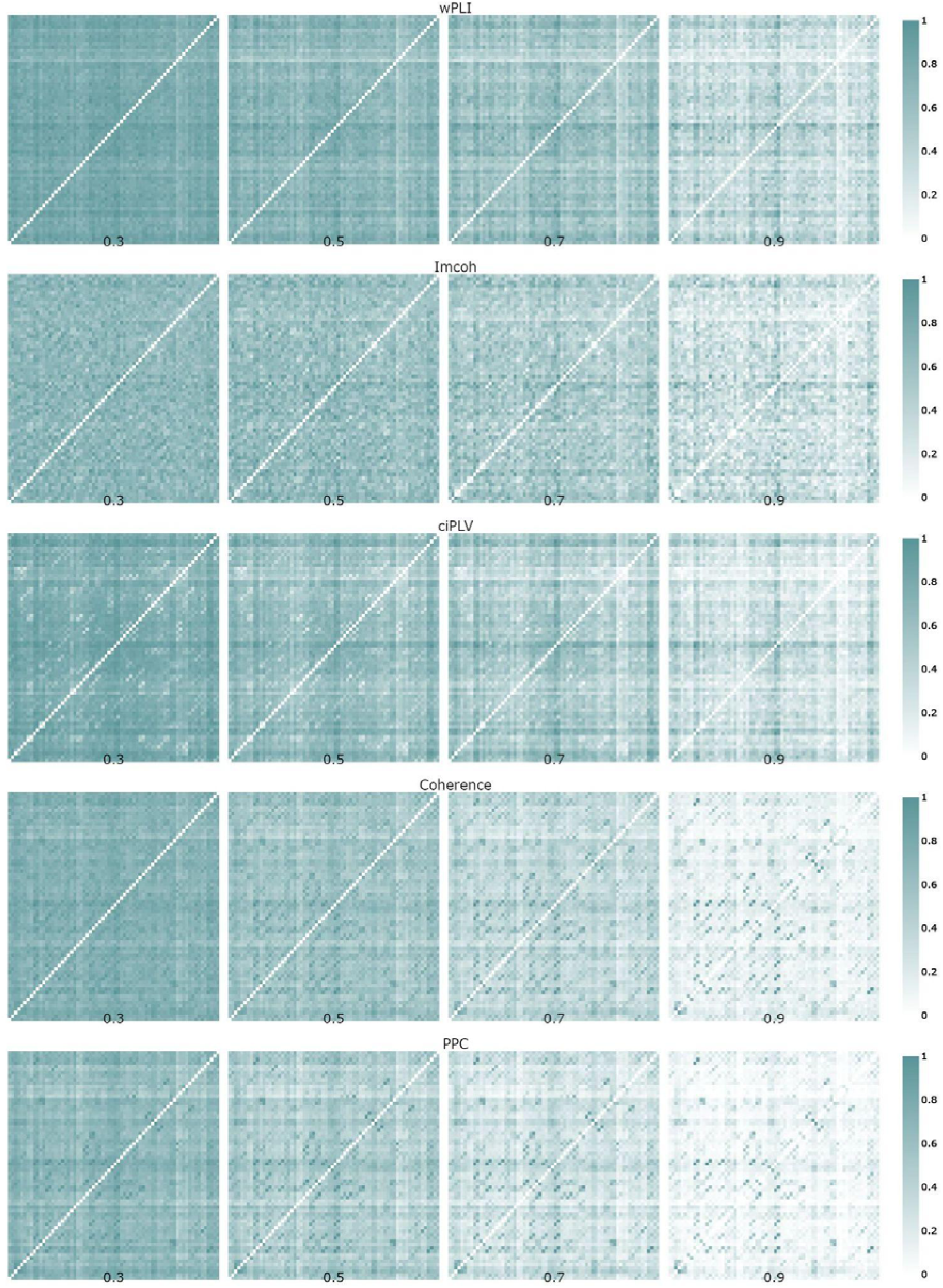
Edge probability graphs across different thresholds in source space. The brightness of cells reflects the relative probability of edge existence. Brighter cells are more frequent in the individual networks in the sample. The edge weights were scaled to the range [0, 1] across all thresholds

The differences in the correlational structure between different thresholds could reveal the linearity and non-linearity in the intra-threshold changes. The highly correlated structure would indicate relative consistency of network topology measures across different thresholds that should indicate more stable networks among the thresholds, while an uncorrelated structure shows instability in the network structure. Thus, we expect a positive correlation matrix with highly correlated adjacent thresholds. We also assume some linear changes in the correlation structure with weaker intra-threshold correlations at high thresholds.

A few points could be derived from our results. First, there are differences in the correlation structure between the different synchronization measures - there are two distinct groups wPLI and ciPLV and imcoh, coherence, and PPC with distinct patterns of correlational structure. Second, Characteristic Path length and clustering coefficient are relatively correlated in adjacent thresholds in all measures. Third, the participation coefficient is not intercorrelated between thresholds at all. Fourth, the strength of intra-threshold correlations tends to decline with higher thresholds, in some cases to negligible levels. Some measures, such as clustering coefficients derived from coherence and PPC, demonstrate negative correlations between measures at different thresholds. Overall, these results show the substantial inconsistency in the network structure across different thresholds and indicate the incongruity between global and local network measures.

### Structure of the functional connectivity graphs on the different thresholds

As the final step in the analysis, to understand the graphs behind the measures, we decided to trace changes in the structure of the functional connectivity graphs via the edge probability graphs.

The edge probability graphs reflect the strongest connections in the connectivity matrices and the stability of connectivity estimation in the sample. If the edge is strong and stable across the sample, the probability of the node existence tends to be 1. Plots 5-6 show that strong connections show up more clearly on the higher thresholds. The thresholding procedure is intended to remove spurious and weak connections in the network and the plots indicate that the probability of the existence of some connections is indeed higher at higher thresholds. However, there seems to be an inconsistency in the structures of individual networks. A thresholding procedure implies that one can expect a ‘core’ of strong connections (and thus a high probability of existence) between nodes persisting across all thresholds with weak connections vanishing. However, the probability of some connections seems to be lowered with the thresholding procedure. For example, relatively strong connections between frontal and parietal/occipital areas estimated with coherence and PPC at the 0.7 threshold are absent at the 0.9 threshold. On the imcoh-derived networks, the probability of connection between electrodes FT9 and P7 weakened at the 0.9 threshold while connections of electrode TP9 are significantly higher at the 0.9 threshold. This inconsistency of scaled probabilities remains in the network analysis of source data e.g. connection between left-hemisphere parahippocampal and right-hemisphere lateral orbitofrontal areas in incoh-derived network is stronger on the 0.5 threshold than on both 0.3 and 0.7 thresholds. This variability of relative probabilities across different thresholds might be a result of various factors. First, it might indicate the inconsistency of the network estimation via proportional thresholding. Second, variability might be a direct consequence of individual differences in brain functional connectivity networks. Third, the variability might be rooted in the very process of EEG functional connectivity estimation, as there seems to be a clear difference between different functional connectivity measures considering the values and topology of networks, but the head-to-head comparison is beyond the scope of this paper.

### Comparison of traditional thresholding approach and data-driven techniques

To avoid the arbitrarily chosen thresholds data-driven approaches have been developed. We chose the orthogonal MST method [17] to represent the data-driven techniques as it is believed that this method creates denser and potentially more meaningful networks. The direct comparison of the networks requires the same density in both networks. The mean density of the OMST-derived graph is 0.45±0.01, which is equivalent to the 0.55 quantile threshold. The following comparison is performed between OMST-derived and 0.55 quantile graphs. The results are presented in Table 2.

**Table 2.**
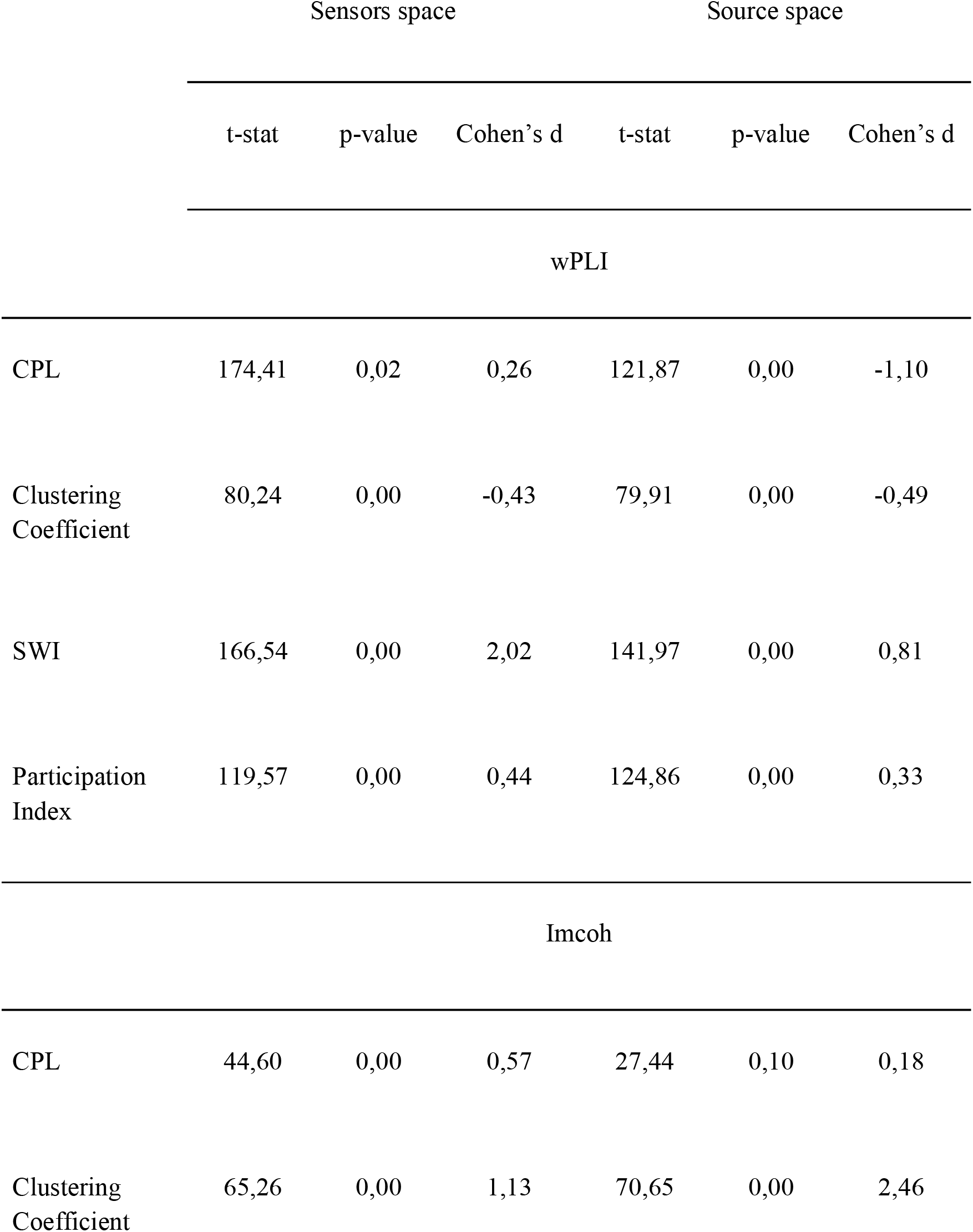
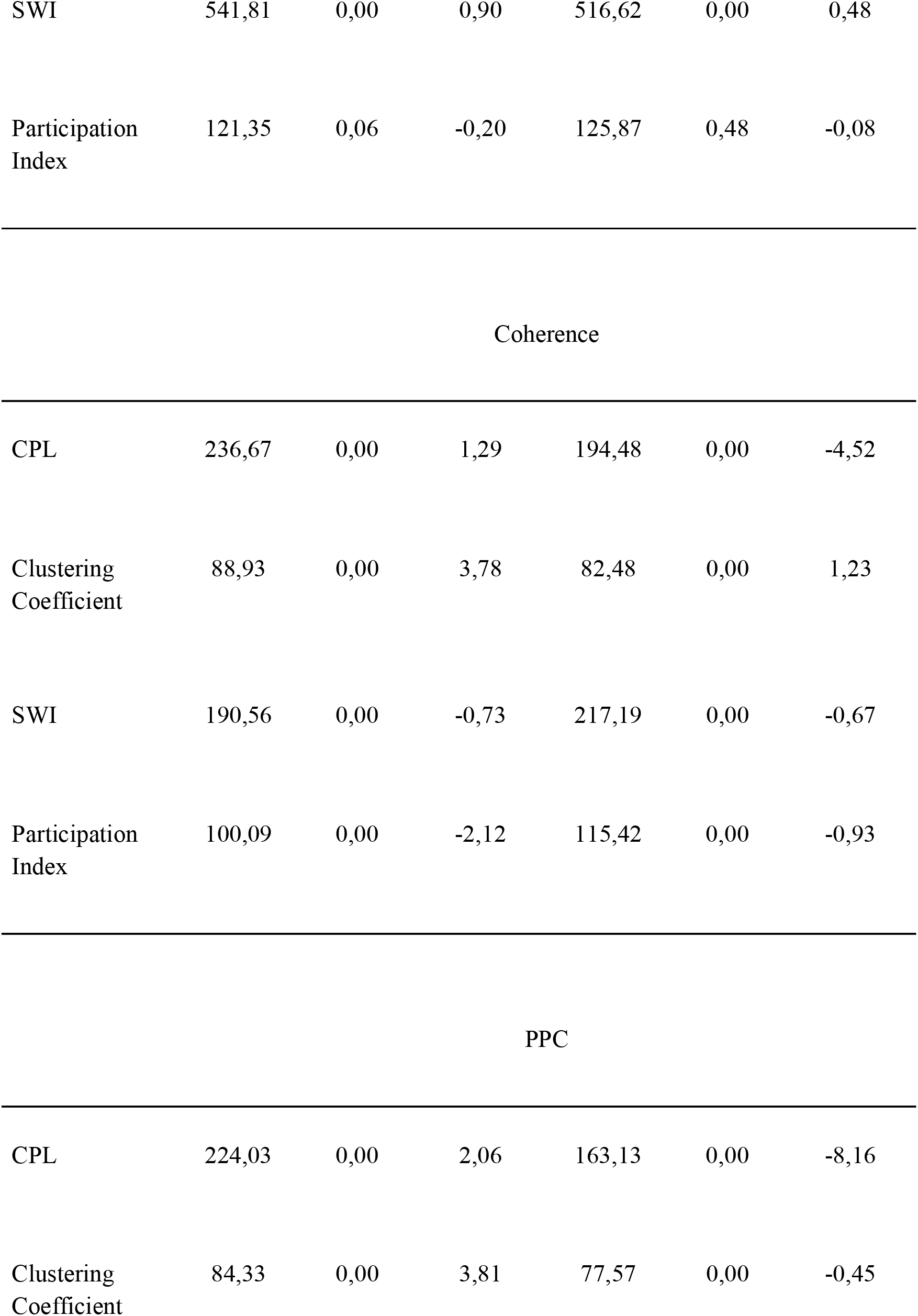
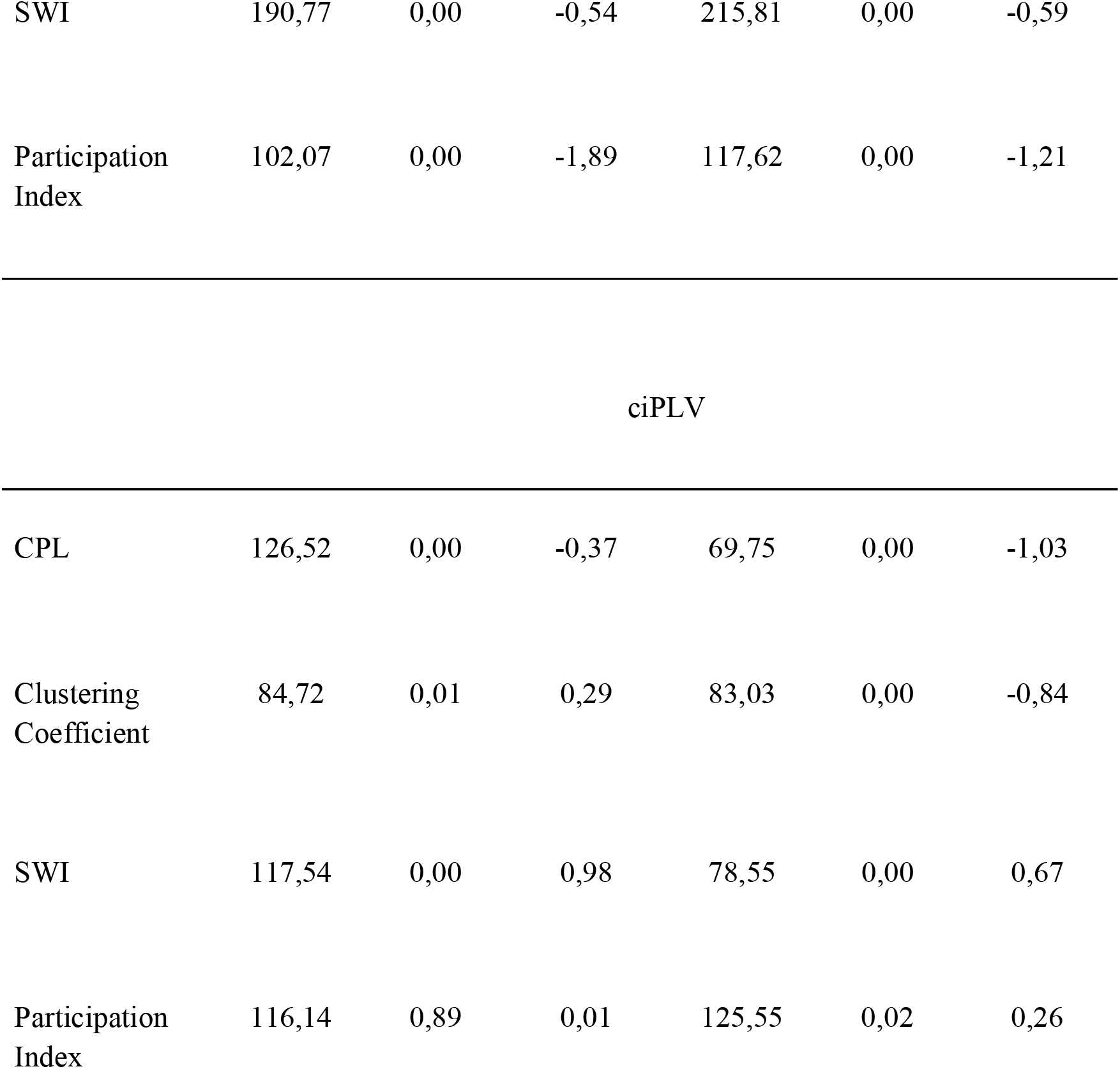
T-statistic comparison between OMST-graph and thresholded graph of similar density

|  | Sensors space |  |  | Source space |  |  |
| --- | --- | --- | --- | --- | --- | --- |
|  | t-stat | p-value | Cohen's d | t-stat | p-value | Cohen's d |
| wPLI |  |  |  |  |  |  |
| CPL | 174,41 | 0,02 | 0,26 | 121,87 | 0,00 | -1,10 |
| Clustering Coefficient | 80,24 | 0,00 | -0,43 | 79,91 | 0,00 | -0,49 |
| SWI | 166,54 | 0,00 | 2,02 | 141,97 | 0,00 | 0,81 |
| Participation Index | 119,57 | 0,00 | 0,44 | 124,86 | 0,00 | 0,33 |
| Imcoh |  |  |  |  |  |  |
| CPL | 44,60 | 0,00 | 0,57 | 27,44 | 0,10 | 0,18 |
| Clustering Coefficient | 65,26 | 0,00 | 1,13 | 70,65 | 0,00 | 2,46 |
| SWI | 541,81 | 0,00 | 0,90 | 516,62 | 0,00 | 0,48 |
| Participation Index | 121,35 | 0,06 | -0,20 | 125,87 | 0,48 | -0,08 |
| Coherence |  |  |  |  |  |  |
| CPL | 236,67 | 0,00 | 1,29 | 194,48 | 0,00 | -4,52 |
| Clustering Coefficient | 88,93 | 0,00 | 3,78 | 82,48 | 0,00 | 1,23 |
| SWI | 190,56 | 0,00 | -0,73 | 217,19 | 0,00 | -0,67 |
| Participation Index | 100,09 | 0,00 | -2,12 | 115,42 | 0,00 | -0,93 |
| PPC |  |  |  |  |  |  |
| CPL | 224,03 | 0,00 | 2,06 | 163,13 | 0,00 | -8,16 |
| Clustering Coefficient | 84,33 | 0,00 | 3,81 | 77,57 | 0,00 | -0,45 |
| SWI | 190,77 | 0,00 | -0,54 | 215,81 | 0,00 | -0,59 |
| Participation<br>Index | 102,07 | 0,00 | -1,89 | 117,62 | 0,00 | -1,21 |
| CPL | 126,52 | 0,00 | -0,37 | 69,75 | 0,00 | -1,03 |
| Clustering<br>Coefficient | 84,72 | 0,01 | 0,29 | 83,03 | 0,00 | -0,84 |
| SWI | 117,54 | 0,00 | 0,98 | 78,55 | 0,00 | 0,67 |
| Participation<br>Index | 116,14 | 0,89 | 0,01 | 125,55 | 0,02 | 0,26 |

The results of comparison between OMST and thresholded graphs show substantial differences between the two. The direction of the changes is ambiguous and unstable across different synchronization measures both in sensor and source spaces, e.g. OMST CPL is lower in PPC - sensor networks and higher in the ciPLV-sensor networks. This might indicate the influence of the individual synchronization strength distribution on the estimation of orthogonal MSTs. In the figures 7-8 we present the probability graphs for the OMST-derived graph and thresholded graph of equal density.

**Figure 7.**
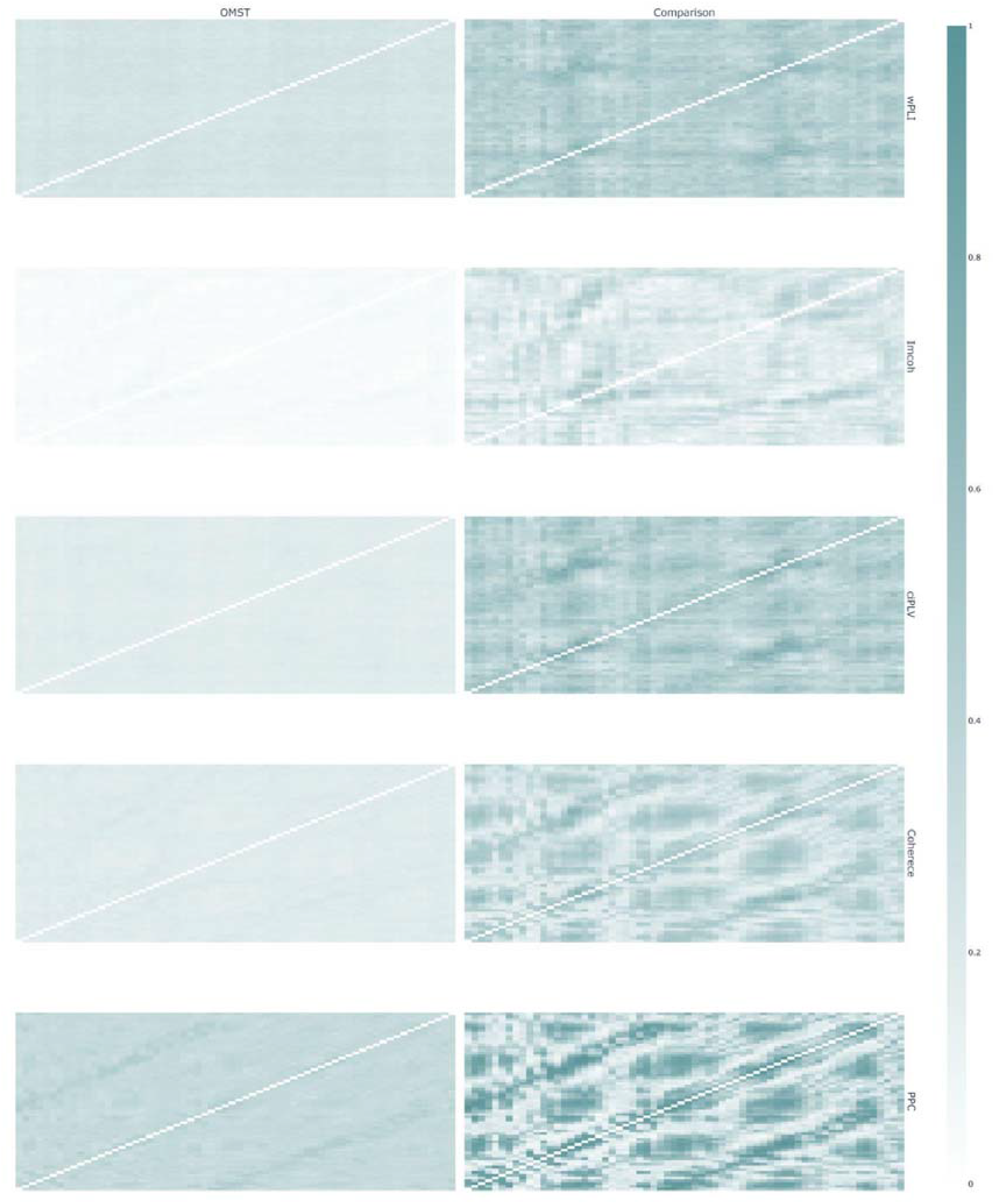
Graph structure for OMST-derived graphs and thresholded graphs of equal density in sensor space. The left column is OMST-derived networks, right - thresholded networks. The brightness of cells reflects the relative probability of edge existence. Brighter cells are more frequent in the individual networks in the sample. Synchronization measures by row: wPLI, Imcoh, ciPLV, coherence, PPC

**Figure 8.**
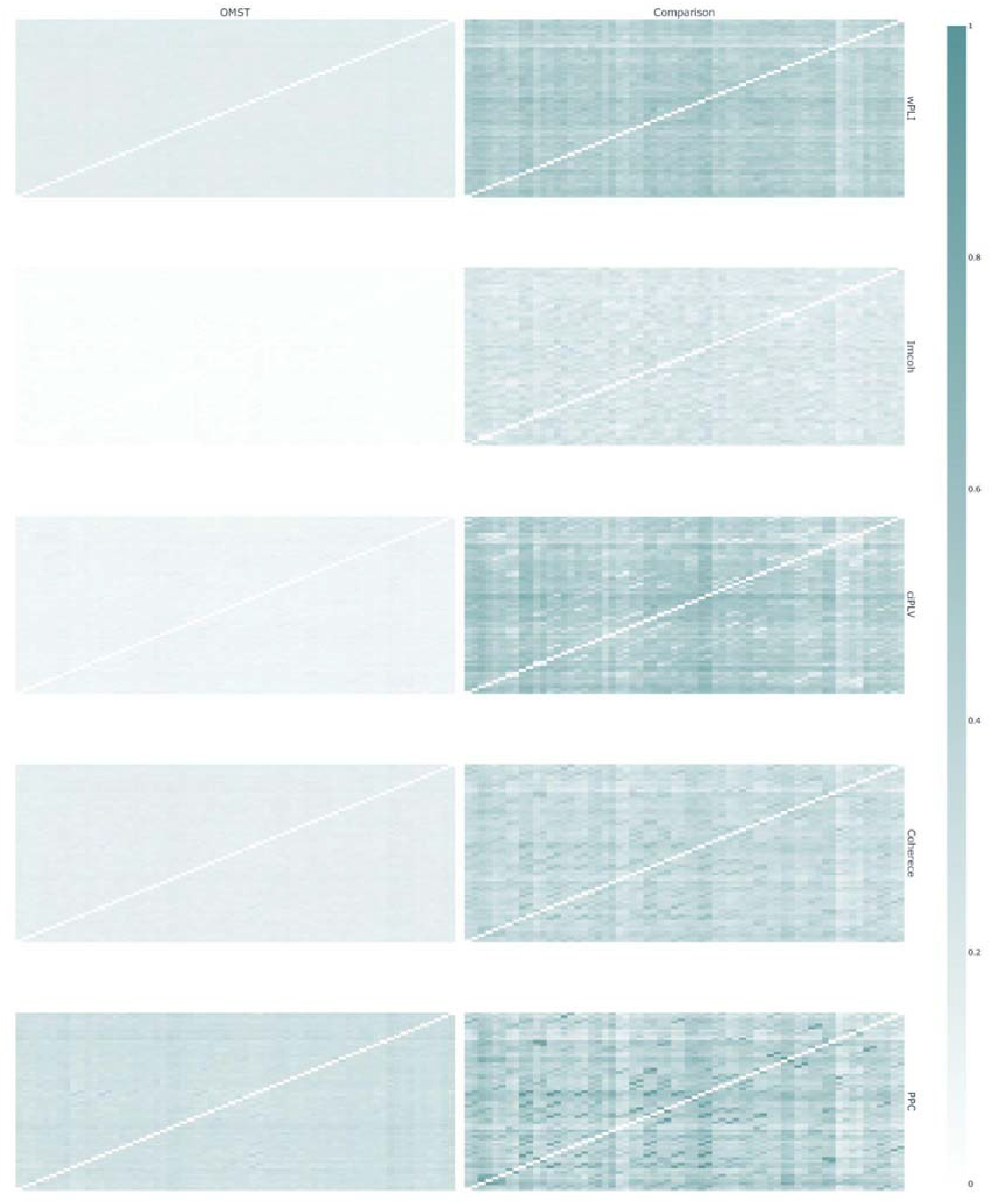
Graph structure for OMST-derived graphs and thresholded graphs of equal density in source space. The left column is OMST-derived networks, right - thresholded networks. The brightness of cells reflects the relative probability of edge existence. Brighter cells are more frequent in the individual networks in the sample. Synchronization measures by row: wPLI, Imcoh, ciPLV, coherence, PPC

In the source space, there is a difference between the edge probability in the OMST-graphs and thresholded graphs. This might reflect individual differences in the graphs. The probability of the edge existence in the OMST-graphs is lower than in the traditional approach and the probability of edge existence is uniformly distributed across all edges in the network. The thresholded networks exhibit areas with higher probability, thus reflecting the existence of more frequent edges in the individual networks across the sample. The structure of OMST-derived probability of edge existence graph might indicate that the OMST algorithm is more prone to individual differences in brain functional connectivity networks.

### Gender differences on different thresholds

The main aim of the functional connectivity studies is to estimate the specificity of brain networks in different conditions, such as gender, different levels of intelligence, or specific task. However, the chosen thresholds could affect the characteristics of the groups, leading to the absence or the presence of the effect of interest depending on the thresholding value. The subjective choice of thresholds results in differences in the absolute values of graph metrics at different levels, values, but these differences will not necessarily be reflected in the experimental results. Here we use the gender differences to model the effect of the thresholding procedure on the outcome of group comparison. The results are shown in figures 9-10.

**Figure 9.**
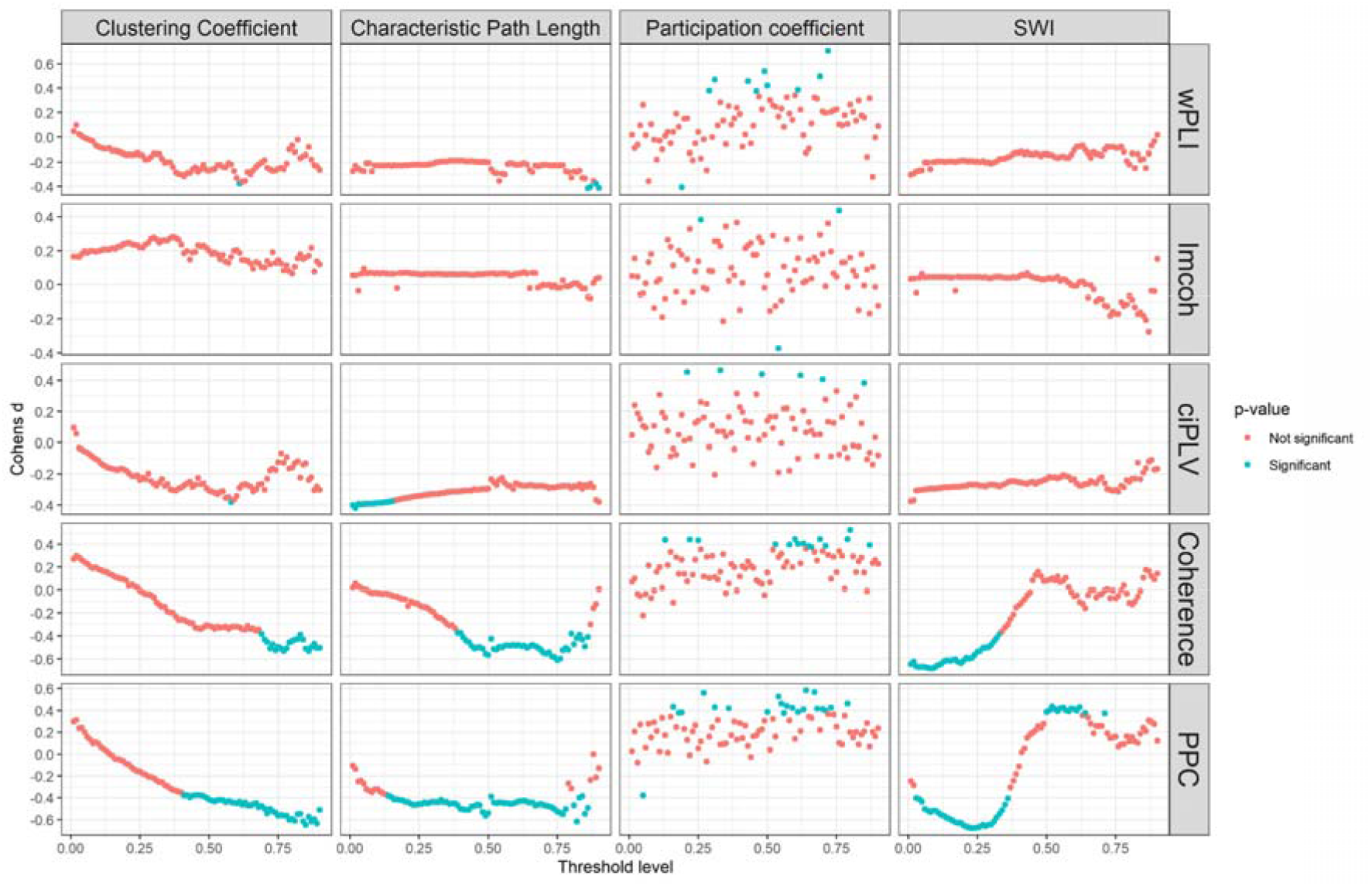
The Cohen’s d of the gender comparison on the different thresholds in sensor space. A permutation t-test was used to estimate the differences between the two groups. The red dots represents the effect as significant (p-value < 0.05), the blue represents the non-significant effect (p-value> 0.05)

**Figure 10.**
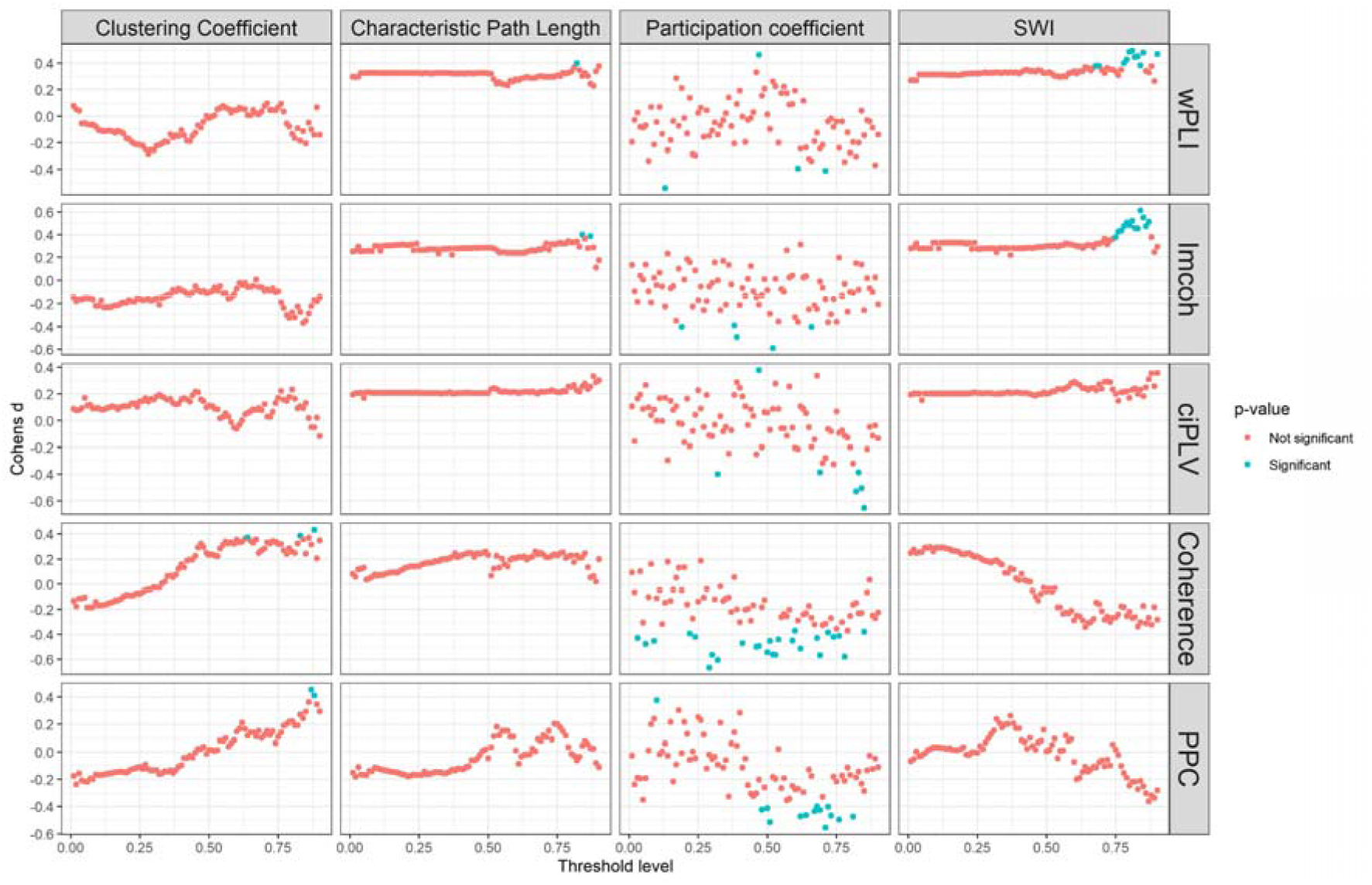
The Cohen’s d of the gender comparison on the different thresholds in source space. A permutation t-test was used to estimate the differences between two groups. The red dots represent the effect as significant (p-value < 0.05), the blue represents the non-significant effect (p-value> 0.05)

As the idea of the thresholding procedure implies the removal of weak or spurious connections in the data, retaining only strong and important ones, one might expect that ‘cleaning’ of the network due to removal on “noisy” connections should give (1) continuous segments of significant differences, (2) significant differences should appear on higher thresholds. Our analysis suggests that our expectations were not fully met, and the results of the group comparisons might differ depending on the threshold level. For example, for the participation coefficient gender differences are non-significant for the majority of the thresholds, but not all of them. It is important to note that those thresholds are scattered across the threshold levels. For other metrics, there are monotonic changes in the magnitude of the group differences with the “significant” intervals. These intervals, however, can start far below the most widely used thresholds, located around the 0.8 threshold. In contrast, at high levels of the thresholds, there is variability in the effect size, which indicates considerable instability in the estimation of group differences. There are also intervals of the significant differences on the lower thresholds (around 0.25 threshold), indicating the possible group differences specifically for the weak connections. Fourth, in our data a peculiar case occurs, the SWI for the PPC in sensor space has two significant intervals with the opposite Cohen’s d values. At low thresholds, the males have higher values of SWI than females, while at high thresholds the effect was the opposite. In this case, the choice of threshold determines not only the conclusion about the presence or absence of an effect but also its direction. Overall, the choice of threshold might significantly alter the experimental conclusions as even two adjacent thresholds might give different results.

## Discussion

The present article aimed to analyze the effect of the thresholding procedure on graph construction from EEG resting state data. According to our results, the thresholding procedure leads to substantial variability in the data. Overall, we have found that:

1. Global graph metrics vary as a function of density level.
2. Density-related graph measures (characteristic path length, clustering coefficient, participation coefficient, and SWI) dynamic have a substantial linear component.
3. Proportional thresholding led to substantial distortions in the graph structure across thresholds, demonstrated by the structure of correlations between different thresholds and the structure of edge probability graphs.
4. Participation coefficient is the metric most affected by the thresholding procedure.
5. Data-driven algorithms may give significantly different results from the traditional thresholding procedure of corresponding density.
6. The chosen threshold may influence the outcome of the analysis (e.g., the presence or absence of the effect of group comparisons).

These discrepancies between the network structure and graph-based metrics on different thresholds are extremely important in light of the widely discussed reproducibility crisis in modern science making it hard to estimate the comparability of the results performed with different analysis pipelines. Overall, according to our results, the way the researcher solves the thresholding issue heavily affects the outcome of the connectivity analysis. Our results add to the previous results by Garrison and colleagues (2015) [13] with fMRI and Civier and colleagues (2019) [16] with DTI, showing that in EEG studies the thresholding should be used with considerable caution. Our study shows that the networks on the different thresholds are obscured by the global measures, as similar values of global measures could be estimated from the variety of structurally different networks. Most importantly, the network structure estimated for the different thresholds may lead to different conclusions of a study. The basic idea of the thresholding procedure (clearing the weak edges out) implies that detecting a stable experimental effect is only possible in the absence of spurious connections (at high thresholds). Our results indicate that the presence of an effect can also be detected at significantly lower threshold levels and its direction can even be the opposite if the weak connections are preserved. This requires a more thoughtful and informed approach to the thresholding procedure.

One of the main challenges in the functional connectivity thresholding problem stems from the fact that there is by now no “ground truth” method on how to define the best threshold value. The data-driven approach based on the MST algorithms has been proposed as a solution to the problem of choosing the arbitrary threshold for network construction [17]. The MST is supposed to reduce the influence of the spurious connections within the brain by considering strong connections within the brain that don’t form cycles or loops thus achieving higher specificity of network topological properties. However, several studies have shown that weak connections can play a significant role when investigating the links between network properties and individual differences in behaviour. The importance of weak connections for network stability has been established for various complex systems, from protein-to-protein interaction [23] to mobile communication networks [24], biological functions [25], or social networks [26]. According to the recent anatomical studies with tracing the connections within the cortical, the brain indeed has a lot of weak connections that have a substantial impact on maintaining the intermodular connections between functional brain modules. Weak connections may also play a significant role in the individual differences in intelligence [27, 28] and its abnormality may be related to the symptomatology of schizophrenic patients [29].

To date, the informed guide on the thresholding procedure seems to be non-existent and many researchers are left to themselves in the procedure that seems crucial for the outcome of the studies. It is still important to provide a basis under the used threshold level and the procedure itself.

Our study was mostly limited to density thresholding, but there are other methods of network extraction, such as component extraction methods, integration measures, windowed thresholding [30], or permutation testing [31] and others that should be tested. Component extraction methods aim at the extraction of a single connected component of the network, e.g. extraction of a Minimum Spanning Tree (MST, a subset of the edges of a connected, edge-weighted undirected graph) or a giant connected component (by removal of weakest connections up to the point when removal of any edge will create an isolated node). Integration approaches require the estimation of measures across various thresholds and the following integration to yield the Area under the curve (AUC) as an integrated measure (REF). Windowed thresholding [30] suggests the separation of an initial network into independent networks by extraction and analysis of networks belonging to discrete windows of edge weights. Permutation testing [31] is based on the assumption that any significant effect should be visible over the continuous range of thresholds. In this approach the significant effects are clustered on the threshold axis, AUC is calculated over the range of thresholds that define clusters and the AUC value is compared to the null model. Recently, an approach to choose the thresholding value according to the entropy of the functional networks [32], as well as some other “optimizational’ measures [33], were suggested, however, its validity and reliability are yet to be established. Integration and permutation methods could be useful when graph measures and experimental effects change monotonously with the change in the thresholding value. However according to our results this is not always the case.. Windowed approaches might struggle with overall integration of the structures of different windowed networks into a one model.

Overall, we believe that there are several ways to improve the analysis of brain functional networks and achieve more robust results. First, the way to overcome the existing thresholding-related problems with network analysis may be to widen the description of the networks with local (nodal) measures and to describe the network topology more thoroughly with different methods, such as graphlets. Graphlet can be defined as an induced subgraph isomorphism class in a graph. Graphlets have been suggested for the analysis of the topological structure of the undirected networks [34], which is important for brain functional connectivity analysis, where networks are mainly undirected by construction. As shown in our study, global network measures, such as average pathlength or clustering coefficient, may be not representative of the topology of the network. As functional connectivity analysis aims to describe connections within the brain, more detailed description and analysis of network topology might be a better option for this purpose than global connectivity measures.

In addition to this, some studies have shown that it is easier to estimate an individual functional connectivity fingerprint than task-related functional connectivity among the group [4, 35]. This is evidence toward the tremendous amount of interindividual variability of network characteristics. More cautious and thorough consideration of individual differences in brain functional networks seems necessary for the better understanding of network properties, underlying psychological functions. be a unique fingerprints of individual brain activity

Third, a possible way to deal with the thresholding problem is linked to the constraining functional connectivity matrices by anatomical structure of connections within the brain. Recently, a two-step pipeline was developed to constrain functional activation with clusters of structural connectivity [36]. However, while the structure-function relationships are an essential part of complex natural systems, it seems that the link between structural connections and functional connectivity can be fundamentally imperfect. Even though the architecture of the structural connectivity networks shapes the boundaries of the functional network, the actual neural dynamics may not be perfectly aligned to it [37].

Fourth, the thresholding problem can be addressed with the null models [38] framework. This framework allows comparison of empirical networks with random or pseudo-random networks with estimating the chance of getting empirical results randomly. Null model network acts pretty much as a null hypothesis for an empirical network to test against. Despite having some methodological considerations [38], null models framework provides a powerful tool in statistical inference. However, we believe that this framework could be improved further. Typical null model networks are created by reshuffling an empirical network or by creation of an artificial network, constrained by properties of the empirical network, such as edge weight distribution or node degree distribution. But imperfections in the very process of functional connectivity estimation could lead to biased networks. We think that spatio-temporal reshuffling of initial biosignals might be a better option for the purpose of creating null models.

Fifth, one might expect more important connections in the network to be more stable over time. Introducing a time dimension and assessing dynamic connectivity during the experiment and choosing only the most stable edges might be valid for the purpose of extracting the most valuable connections in networks.

Last, we think that the introduced concept of edge probability graphs might be useful in estimation of the most important edges in the networks. The edge probability graph removes the effect of variable absolute connectivity strength in the sample, retaining the position of the node strength among the other edges in the individual network. That could ease the non-stationarity of brain functional networks, caused by individual differences and imperfections in the process of connectivity estimation. Assessing the most probable edges instead of the strongest connections or averaging across multiple adjacency matrices might be a better tool in description and analysis of diverse individual networks, network stability in time or trial-by-trial stability and variability. However, this approach should be developed further in more detail.

The variability in the ways of graph-metrics estimation related to thresholding procedure is just a part of a bigger problem of the variability in the analysis pipelines in neuroimaging data. Recently, the results from 70 research teams focused on analyzing the same datasets showed that the analytic choices, made by researchers, lead to the substantial affect variability in the research conclusions [39]. To directly estimate the role of such variability in the EEG research the open EEG Many Pipelines has been launched (https://www.eegmanypipelines.org/).

What could be done to choose among the vast amount of possible analytic choices? One solution may lie in data sharing, that opens the possibility to test whether different analysis will lead to the comparable results (for the discussion regarding open data practices see [40]). Another way to overcome the huge variability in the data analysis are pre-registration practices [41]. The idea of pre-registration is to have a record (public or stored in the journal editorial office) of the detailed analysis plan prior to the data collection. Current experience shows that pre-registration leads to higher level of the replicability of the research [42]. Finally, analysis of complex datasets can be performed according to the multiverse approach [43]. According to the multiverse approach the overall conclusion regarding the effects of interest can rely only on comparing various analysis procedures [44]. To additionally increase the integrity and the reproducibility of these analyses the analysts can also be “blinded” regarding the initial hypothesis of the research. Recently, the guideline for multi-analyst research emerged [45] wrapping up the techniques of strengthening the robustness of scientific results. As the different analysis pipelines used by research groups could possibly lead to different conclusions, authors encourage independent researchers to perform an analysis on the same data, inferring the robust result where results from the different researchers converge.

To sum up, in the present study we have shown that the thresholding procedure might be crucial for the outcome of the experiment. While the threshold level choice is arbitrary and is unsupported by compelling reasons, this procedure is an additional source of uncertainty in the experimental results in an already indefinite field. In agreement with the broader problem of the analytic choices in neuroimaging studies, the analysis of the brain connectivity can be highly dependent on choosing the thresholds for connectivity matrix construction with no best way to choose between different analytic options. Different approaches, such as multiverse analysis or analysis by different research teams are advocated for.

## Methods

### Participants

164 participants, aged from 17 to 34 years (M=21.7, SD = 3.36, 30% female) with no report of head traumas or psychiatric or neurological disorders took part in our study. Written informed consent was obtained from the participants or their legal guardians for their participation in the study. No monetary reward was given to the participants. The study was conducted according to the guidelines of the Declaration of Helsinki and approved by the ethical committee of the Psychological Institute of Russian Academy of Education.

### Experimental procedure and data acquisition

Brain Products ActiChamp was used for the 64-channel EEG recording with a 500Hz sampling rate. Cz electrode was used as a reference. EEG preprocessing included four steps: 1) Downsampling to 256Hz, 2) Re-reference to a common reference, 3) Filtering from 1Hz to 30Hz and 4) artefact removal. An artifact removal consisted of manual removal of artifacts, removal of ocular artifacts via an infomax Independent Component Analysis and topographic interpolation of the noisy channels.

The experimental procedure consists of five alternating 2-minute intervals with closed and open eyes. Total 10 minutes per participant were recorded with 3 closed-eyes intervals. Only closed-eyes condition was used in the study.

### Source reconstruction

Source reconstruction was performed using the pipeline mentioned in the documentation to the MNE package [46]. The source reconstruction was performed based on a three-layer BEM model with 1026 sources per hemisphere. The three layers represent the inner skull, outer skull and outer skin and the conductivity of the layers was standard for the MNE package (0.3, 0.006 and 0.3). To construct the BEM model, the standard FreeSurfer model head, Colin27, was used. The individual source reconstruction was performed with the individual inverse operator using the dSPM method. Cortical parcellation was achieved via the MNE function mne.extract_label_time_course with the “pca_flip” method with the Desikan-Killiany Atlas [47].

### Estimation of synchronization

Five measures were used to estimate synchronization: weighted phase lag index (wPLI [48]), Imaginary part of Coherency (imcoh, [49]), coherence, corrected imaginary phase locking value (ciPLV, [50]) and the Pairwise Phase Consistency (PPC, [51]). Each of these measures has their own assumptions and are easily available via common EEG-analysis packages (e.g. MNE for Python). Synchronization was estimated in alpha band 8-13 Hz frequency range for sensor and source space with each synchronization measure for eyes-closed condition.

### Graph analysis

The NetworkX package [52] was used for the graph analysis. Each connectivity matrix was considered as a weighted graph adjacency matrix. Threshold values started from 0.01 quantile of the matrix weights distribution (99% of the connections included) to 0.99 (1% of the strongest connections included) with 0.01 step. Edges with weight below the threshold were eliminated. On each step, disconnected nodes were removed and are reported as isolated nodes. In case the graph divides into separate subgraphs only the biggest component remains. The graph structure on each threshold was constructed based on the probability of edge existence in the whole sample. The probability of edge existence for each edge was defined as the ratio of adjacency matrices with non-zero edge weight to the total number of adjacency matrices. The probability graphs were constructed for each threshold. The probabilities of edge existence for each measure were scaled across all thresholds to [0:1] range for better visibility of edges with high probability.

Another way to construct the graph is based on the minimum spanning tree (MST) algorithms. The minimum spanning tree is a set of edges that connects all nodes with minimum possible edge weight. We use the orthogonal MST algorithm developed by G. Dimitriadis [17]. The orthogonal MST method implies the consequent extraction of MSTs from the full graph. The MST on each step is removed from the original graph (all edges belonging to the MST are set to zero), but it is transferred to a new, thresholded network. The algorithm ends when an equilibrium between global efficiency and edge removal cost is reached. We compare the data-driven graphs with the same density traditionally thresholded graph.

For each graph, we calculated four measures to describe the network structure: Characteristic Path Length, Clustering Coefficient, participation coefficient and SWI.

*The Characteristic path length* is defined as the median of the distribution number of edges between two nodes in the shortest distance between them.

*The clustering coefficient* is a measure of the tendency of nodes to form interconnected groups. For each node clustering coefficient is a proportion of the existing number of connections in a neighbourhood to a total number of possible connections.

*Participation coefficient* is a measure of node edge distribution across networks calculated for each node. Nodes with uniform link distribution with other nodes will have participation coefficient 1, if a node has edges among one specific community, the participation coefficient will be 0. The mean participation centrality across all nodes is presented.

SWI (Small-world Index, [53]) is a measure of a small-worldness of the graph via the comparison of the characteristic path length and clustering coefficient of a given graph with the measures in the set of random graphs.

### Statistical analysis

Linear regression analysis was performed to estimate the graph measure dynamic. A permutation t-test was used to estimate the differences between traditional and data-driven approaches and group differences. Spearman correlation coefficient was used for estimation of the similarity between different thresholds. The connectivity matrices were created via the MNE package [46, 54], graph analysis was performed in NetworkX [47], OMST-graphs were created via authors package in Matlab (https://github.com/stdimitr/multi-group-analysis-OMST-GDD), statistical analysis was performed in R in stats (https://www.r-project.org/) and exactRankTests (https://CRAN.R-project.org/package=exactRankTests) packages

## Authors Contribution

T.A.: Conceptualization, Methodology, Software, Formal analysis, Visualization, Writing – Original Draft; I.Z.: Conceptualization, Methodology, Resources, Writing – Original Draft, Writing - Review & Editing; A.T.: Investigation, Data Curation, Writing - Review & Editing; S.M.: Writing - Review & Editing, Supervision, Project administration

The author(s) declare no competing interests.

The datasets generated during and/or analysed during the current study are available from the corresponding author on reasonable request.

## Notes

### Competing Interest Statement

The authors have declared no competing interest.

### Summary of Updates

Few grammar mistakes were corrected

